# Covert actions of epidural stimulation on spinal locomotor circuits

**DOI:** 10.1101/2024.06.18.599598

**Authors:** D. Leonardo Garcia-Ramirez, Jenna R. McGrath, Ngoc T. Ha, Jaimena H. Wheel, Sebastian J. Atoche, Lihua Yao, Nicholas J. Stachowski, Simon F. Giszter, Kimberly J. Dougherty

## Abstract

Spinal circuitry produces the rhythm and patterning of locomotion. However, both descending and sensory inputs are required to initiate and adapt locomotion to the environment. Spinal cord injury (SCI) disrupts descending controls of the spinal cord, producing paralysis. Epidural stimulation (ES) is a promising clinical therapy for motor control recovery and is capable of reactivating the lumbar spinal locomotor networks, yet little is known about the effects of ES on locomotor neurons. Previously, we found that both sensory afferent pathways and serotonin exert mixed excitatory and inhibitory actions on lumbar interneurons involved in the generation of the locomotor rhythm, identified by the transcription factor Shox2. However, after chronic complete SCI, sensory afferent inputs to Shox2 interneurons become almost exclusively excitatory and Shox2 interneurons are supersensitive to serotonin. Here, we investigated the effects of ES on these SCI-induced changes. Inhibitory input from sensory pathways to Shox2 interneurons was maintained and serotonin supersensitivity was not observed in SCI mice that received daily sub-motor threshold ES. Interestingly, the effects of ES were maintained for at least three weeks after the ES was discontinued. In contrast, the effects of ES were not observed in Shox2 interneurons from mice that received ES after the establishment of the SCI-induced changes. Our results demonstrate mechanistic actions of ES at the level of identified spinal locomotor circuit neurons and the effectiveness of early treatment with ES on preservation of spinal locomotor circuitry after SCI, suggesting possible therapeutic benefits prior to the onset of motor rehabilitation.

## INTRODUCTION

Lumbar spinal neurons that generate locomotor rhythm and pattern require supraspinal descending and sensory afferent inputs to initiate and adapt locomotion to the environment (*1–5*). Spinal cord injury (SCI) disrupts the descending control of the spinal locomotor circuitry and often results in paralysis (*6–8*). After SCI, sensory afferent feedback becomes crucial as a principal source of input to modulate spinal locomotor networks (*7, 9–11*). In fact, abolishment of afferent feedback, or disruption of interneurons interposed between afferents and locomotor circuit neurons, prevents locomotor recovery after SCI (*12–15*). Strategies to regain locomotor control after SCI, such as epidural stimulation (ES), aim to reactivate dormant locomotor circuitry by stimulating afferent fibers (*8, 16–21*). However, the actions of ES on spinal locomotor interneurons are still largely unknown.

After SCI and following ES, there are changes that affect the spared spinal circuits (*19, 22–25*). Studies focused on genetically-defined neuronal types have revealed differential mechanisms of SCI-induced plasticity at the level of synaptic input connectivity, intrinsic cellular excitability, and neurotransmitter phenotype (*14, 22, 25–29*). Neurons expressing the transcription factor Shox2 have been implicated in the generation of the locomotor rhythm and pattern (*30*) and, thus, represent a logical target for post-SCI therapeutics. We previously showed that the excitability and intrinsic properties of Shox2 interneurons are unaltered at chronic timepoints after a complete spinal thoracic (T8/9) transection (*25*). However, there were significant changes to the sensory input pathways to and serotonergic modulation of Shox2 interneurons with chronic SCI, both biasing toward excitatory effects (*25*).

Here, to elucidate the effects of ES at the level of the Shox2 interneurons, we performed electrophysiological recordings from Shox2 interneurons in lumbar spinal slices from chronically complete transected mice, a subset of which received daily sub-motor threshold ES. We found that the sensory afferent input to Shox2 interneurons that is mainly excitatory in the SCI mice, is heterogeneously excitatory and inhibitory in SCI mice that received ES. ES, however, does not change the intrinsic excitability of Shox2 interneurons. Similarly, the serotonin supersensitivity, mediated by 5-HT_2B/2C_ receptors, observed in Shox2 interneurons from SCI mice is not present in Shox2 interneurons from SCI mice that received ES. Additionally, we found that ES-induced circuit effects remain even after discontinuing ES, but starting ES after SCI-induced plasticity has occurred is ineffective at reversing the changes. Overall, these data suggest that ES prevents, but does not reverse, SCI-induced plasticity at the level of the Shox2 interneurons.

## RESULTS

### Sub-motor threshold epidural stimulation does not improve locomotor function

One of the goals of epidural stimulation (ES) is to activate locomotor spinal neurons to restore motor function after SCI. ES in combination with other rehabilitation strategies such as treadmill training and/or activation of spinal circuitry with serotonin receptor agonists has led to locomotor improvements in humans with spinal cord injury and animal models with incomplete SCI (*8, 10, 18, 28, 31–33*). In order to test the effects of epidural stimulation (ES) on locomotor function and various electrophysiological parameters at the level of lumbar Shox2 interneurons, mice received a complete spinal transection at thoracic (T) 8/9 level and a subgroup of mice were implanted with ES electrodes over the lumbar spinal segments during the same surgery (SCI+ES group). ES was administered starting one week after surgery and continued for five days per week until the day prior to the slice electrophysiology experiment (Figure 1A and 1B, see Methods). In initial trials, we found that ES above motor threshold produces co-contraction of hindlimb muscles, likely due to the size of the mouse cord and activation of the ventral roots (*9, 16*). Thus, we reduced the stimulation intensity to be below the threshold for muscle twitch (sub-motor threshold).

**Figure 1.**
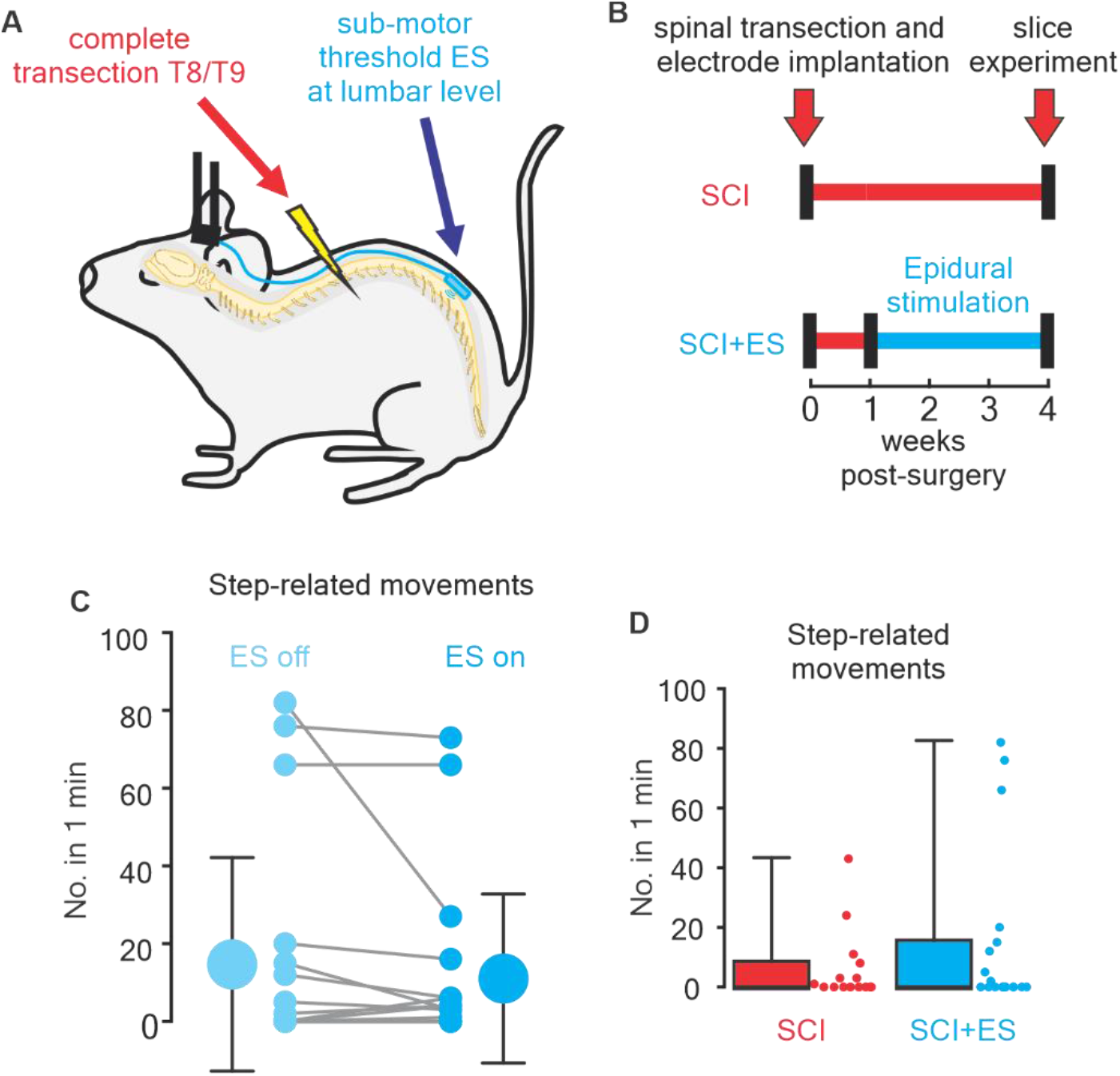
Timeline and locomotor effects of ES. **(A)** Schematic of the complete transection spinal at T8/T9 and implantation of electrodes for epidural stimulation (ES) at lumbar level. **(B)** Experimental timeline for SCI (red) and SCI+ES (cyan) mice. **(C)** Direct comparison of the number of step-related movements in one minute with ES off (left, light cyan) and ES on (right, cyan). The large dots indicate mean and standard deviation. **(D)** Comparison of the number of step-related movements in one minute from SCI (red) and SCI+ES (cyan) mice during the week of slice experiment. Here, the SCI+ES mice values were exclusive to measurements taken with ES off.

In order to identify locomotor improvements induced by ES acutely and over a chronic timeframe, in a subset of SCI+ES (N=19) mice, we measured the number of step-related movements. The step-related movements included mainly ankle flexions and knee flexions but, in some cases, dorsal steps and plantar steps were produced. We tested the acute effects of sub-locomotor threshold ES by comparing the locomotor performance of a subset of SCI+ES mice with ES turned off (ES off) and turned on (ES on). There was no change in step-related movements (ES off: 14.6 ± 27, ES on: 11.0 ± 22; Wilcoxon matched-pairs test, W=-22, p=0.3; Figure 1C). Although acute enhancements in locomotor function were not apparent, we hypothesized that chronic activation may result in enhanced locomotor-like movements (*10, 34*). Therefore, we quantified and compared the step-related movements in SCI+ES mice while the ES was off to SCI mice (N=15) at the same 4 week post-SCI time point. Most of the mice in both groups did not display any step-related movement (55.9%, 19 of 34 mice). In mice with hindlimb movements, variability was high and there were no significant differences in the number of step-related movements (SCI mice: 6.2 ± 12.1, SCI+ES mice: 14.6 ± 27.5; Mann-Whitney test, U = 137, p = 0.8; Figure 1D). These results show that sub-motor ES does not improve locomotion in our complete transection model.

These results motivated us to ask whether prolonging ES treatment and enhancing sensory input by delivering ES while the mice are on a moving treadmill would lead to locomotor improvements. Therefore, in another set of mice, we doubled the length of time of the ES to 6+ weeks and delivered it while the mice were harnessed over a moving treadmill (SCI+ES+tr^long^, Supplementary Figure 1A). We included control groups of mice that received ES in the cage for the same time course (SCI+ES^long^) as well as mice that were on the treadmill without ES (SCI+tr^long^). We found no differences in the number of step-related movements (Supplementary Figure 1B). Thus, the lack of motor improvement with sub-motor threshold ES in our model allows for the determination of plasticity that results from chronic ES without input or influences of observable motor gains in function.

### Sensory afferent input to Shox2 interneurons is altered by ES

SCI-induced plasticity includes changes in sensory afferent transmission to spinal neurons (*35–37*) and has been shown for afferent pathways to Shox2 interneurons (*25*). After SCI, ES is used as an attempt to regain locomotor control via activation of afferent fibers (*9, 16*). Here, to identify potential effects of ES on the sensory afferent pathways to Shox2 interneurons after SCI, we performed whole cell patch clamp recordings on visually identified Shox2 interneurons from both SCI and SCI+ES mice 4.3 ± 0.5 weeks after surgery (Figure 1). We recorded postsynaptic potentials evoked by dorsal root stimulation in Shox2 interneurons from SCI and SCI+ES mice (Figure 2A) in transverse slices with dorsal rootlets attached. The percentage of tested Shox2 interneurons receiving any postsynaptic potential from sensory afferent stimulation was similar in both SCI mice and SCI+ES mice, 20% (12 of 60) and 13% (9 of 68) neurons, respectively (binomial t-test two-tailed, p = 0.22). This shows that Shox2 interneurons from SCI and SCI+ES mice receive sensory afferent inputs in similar proportions.

**Figure 2.**
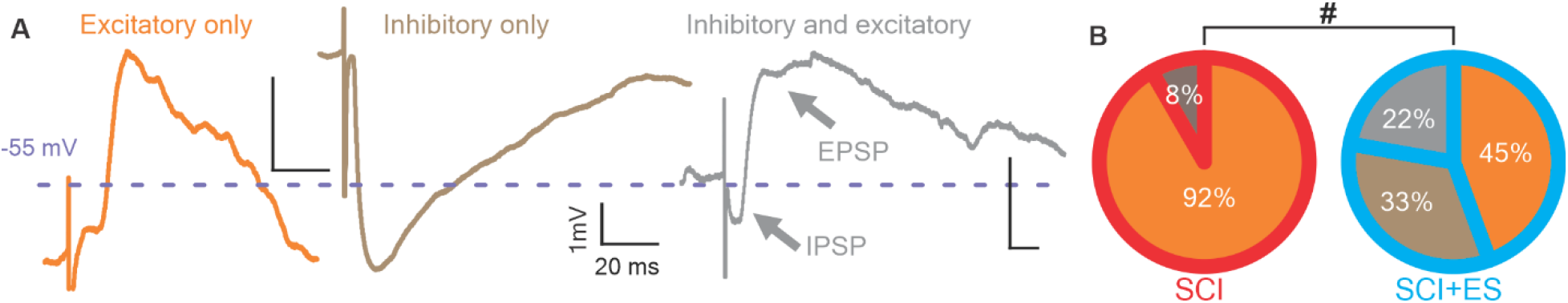
Primary afferent-evoked inputs to Shox2 interneurons from SCI and SCI+ES mice. **(A)** Examples of Shox2 interneurons that receive excitatory (EPSP, orange trace), inhibitory (IPSP, brown trace), and mixed (both IPSP and EPSP, gray trace) synaptic inputs following the stimulation of dorsal roots. **(B)** Incidence of input type received by Shox2 interneurons from SCI and SCI+ES mice in response to sensory stimulation. Excitatory only (orange), inhibitory only (brown), and mixed excitatory and inhibitory (gray). The incidence of type of input to Shox2 interneurons was significantly different between SCI and SCI+ES mice (chi-square test, p=0.02).

Considering only Shox2 interneurons with input from dorsal root stimulation, we subclassified the individual interneurons as receiving only excitatory inputs (Figure 2A, orange trace), only inhibitory inputs (Figure 2A, brown trace), or mixed inhibitory and excitatory inputs (Figure 2A, gray trace). In most cases of mixed inputs, the inhibitory postsynaptic potential was the shortest latency response, with an excitatory postsynaptic potential cutting the inhibitory potential short (as in Figure 2A, gray). Shox2 interneurons from SCI mice received mostly excitatory synaptic inputs (92%, 11 of 12 neurons) with only 1 neuron receiving inhibitory inputs (Figure 2B). In contrast, only 45% (4 of 9) of Shox2 interneurons from SCI+ES mice received exclusively excitatory inputs. Of the remaining neurons, 55% (5 of 9) responded with either inhibitory (3 of 9) or mixed (2 of 9) inputs (Figure 2B). These data show that the afferent input pathways to Shox2 interneurons in the SCI+ES mice are significantly different from the untreated SCI group (*χ*^2^(_2_)=7.3, p=0.02), suggesting that ES alters the SCI-induced plasticity of sensory afferent pathways to Shox2 interneurons.

Afferent input to Shox2 neurons was also tested in slices from mice that received ES on the treadmill over a longer time course and the associated controls. Inputs to Shox2 interneurons in SCI+tr^long^ mice were almost entirely excitatory (86%, 6 of 7, Supplementary Figure 2A), similar to SCI mice. In contrast, like the SCI+ES group, inhibitory inputs to Shox2 interneurons were observed in SCI+ES^long^ (38%, 3 of 8 exclusively inhibitory; 13%, 1 of 8 mixed) and SCI+ES+tr^long^ (55%, 6 of 11 exclusively inhibitory; 18%, 2 of 11 mixed) mice. There were no differences in the types of inputs when comparing SCI+ES^long^ with SCI+ES+tr^long^ mice (*χ*^2^ =2.1, p=0.4) or in the inputs received in SCI+ES^long^ and SCI+ES mice (*χ*^2^ =0.8, p=0.7). Taken together, the results demonstrate that, despite the lack of motor improvements, sub-motor threshold ES leads to changes in afferent input pathways to Shox2 interneurons.

### Intrinsic properties of Shox2 interneurons are not altered by sub-motor threshold ES

The intrinsic excitability of Shox2 interneurons may affect their responses to afferent input. Thus, to elucidate possible differences in the passive and active properties of Shox2 interneurons between SCI and SCI+ES mice, we measured and compared the intrinsic properties of Shox2 interneurons from both SCI (n=60) and SCI+ES (n=49) mice. (Figure 3 and Table 1). We did not find significant differences in the resting membrane potential, input resistance, time constant, rheobase current and action potential threshold. This demonstrates that sub-motor threshold ES does not affect the intrinsic properties and the excitability of Shox2 interneurons compared to SCI alone.

**Figure 3.**
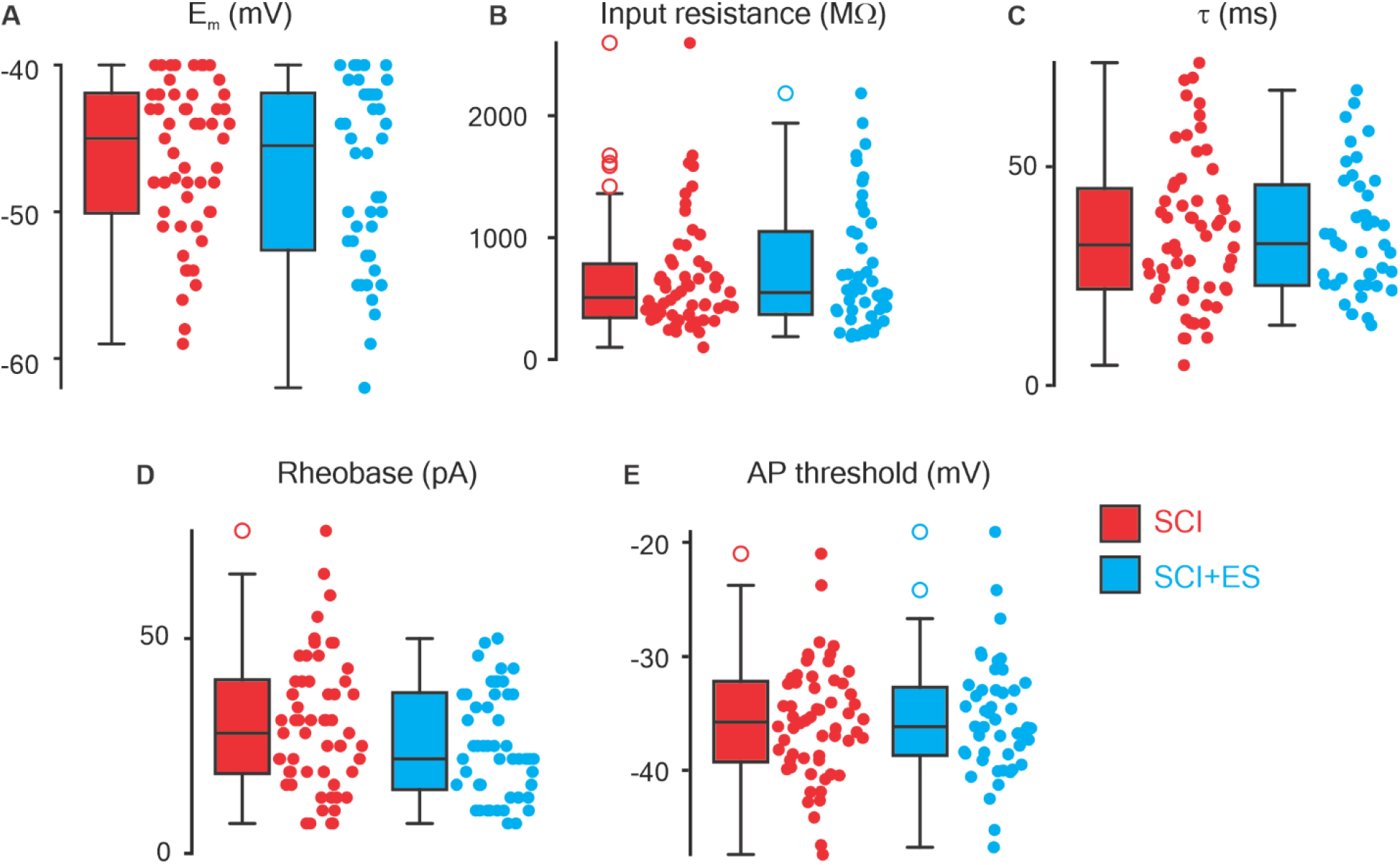
Intrinsic properties of Shox2 interneurons from SCI and SCI+ES mice. Comparisons of intrinsic properties of Shox2 interneurons from SCI (red) and SCI+ES (cyan) mice revealed no significant differences in **(A)** resting membrane potential (E_m_), **(B)** input resistance (R_in_), **(C)** time constant (τ), **(D)** rheobase or **(E)** action potential threshold.

**Table 1.** Comparison of intrinsic properties between Shox2 interneurons from SCI and SCI+ES mice.

| | SCI<br>(mean ± SD) | SCI+ES<br>(mean ± SD) | unpaired *t* test<br>or Mann-Whitney | *p* |
| --- | --- | --- | --- | --- |
| Membrane potential (mV) | -46.2 ± 5 | -47.3 ± 6 | U=1145 | 0.5 |
| Input resistance (MΩ) | 654.0 ± 425 | 732.3 ± 511 | U=1382 | 0.6 |
| Time constant (ms) | 35.0 ± 14 | 35.0 ± 17 | U=1216 | 0.9 |
| Rheobase (pA) | 29.9 ± 15 | 25.2 ± 15 | U=1214 | 0.1 |
| Action potential threshold (mV) | -35.8 ± 5 | -35.4 ± 5 | *t*<sub>(107)</sub>=0.3 | 0.7 |

### Serotonergic modulation of Shox2 interneurons

SCI disrupts the supraspinal serotoninergic modulation of spinal neurons, resulting in plasticity at the level of the expression of receptors on their spinal neuronal targets, including Shox2 interneurons and motoneurons (*25, 27, 38–43*). To identify whether ES affects the serotonergic modulation of Shox2 interneurons, we applied two concentrations of serotonin during our recordings: 0.1 µM (low serotonin) and 10 µM (high serotonin). These were selected based on our prior work which demonstrated that Shox2 interneurons from uninjured adult mice show an overall hyperpolarizing response to 0.1 µM serotonin and depolarize to 10 µM serotonin (*25*). To determine the serotonergic modulation of the excitability of Shox2 interneurons, we considered the differences between the pre- and post-drug steady state for the resting membrane potential (Figure 4A), rheobase, action potential threshold, spontaneous action potential firing frequency during a continuous recording (Figure 4B), and action potential frequency in response to a depolarizing current step (Figure 4C, Table 2, and Supplementary Table 1). To include these five parameters as a single descriptor of changes in the excitability after serotonin, we performed principal component analysis on the differences between the pre- and post-drug steady of the variables to reduce dimensionality. We found that the first principal component (PC1) represents 39.6% and the second principal component (PC2) represents 20.7% of the total variability (Table 3). The loadings of each of the five electrophysiological parameters included within the PCA were range from 0.25 to 0.59 in PC1, suggesting that all variables contribute similarly to PC1 (Supplementary Table 2). The pre-drug conditions of each cell projection onto PC1 and PC2 were then set to zero, and positive values represent increases in excitability, while negative values indicate reductions in excitability in response to serotonin. Shox2 interneurons from SCI mice displayed positive PC1 values of z-score (increased excitability), for both low (0.4 ± 0.5) and high (0.7 ± 1.3) concentrations of serotonin (Figure 4D, 0.1 µM: one-sample *t* test, *t*(_13_) = 2.6, *p* = 0.02; light red. 10 µM: one-sample *t* test, *t*(_9_) = 1.7, *p* = 0. 1, red). Interestingly, low concentrations of serotonin reduced the excitability of Shox2 interneurons from SCI+ES mice (PC1 z-score = −1.5 ± 0.7, one-sample *t* test, *t*(_9_) = 6.8, *p* = 0.0001; Figure 4D, light cyan) and excitability was not significantly changed by high serotonin concentrations (PC1 z-score = −0.7 ± 1.9, one-sample *t* test, *t*(_7_) = 1.1, *p* = 0.3; Figure 4D, cyan). PC2 values were not different among both groups. The response to low concentrations of serotonin was significantly different between Shox2 interneurons from SCI and SCI+ES mice (unpaired *t* test, *t*(_22_) = 7.5, *p* = 0.0001; Figure 4D, Table 3). Shox2 interneuron excitability changes in response to low concentrations of serotonin in SCI+tr^long^ mice were similar to SCI mice (Supplementary Figure 2, Table 7; unpaired *t* test, *t*(_24_) = 2.0, *p* = 0.06) and, similarly to the 3-week SCI+ES mice, Shox2 interneurons from SCI+ES^long^ and SCI+ES+tr^long^ mice did not display the excitatory response to low concentrations of serotonin (Supplementary Figure 2B, Table 7). These data demonstrate that ES modifies the responses of Shox2 interneurons to serotonin observed after SCI, eliminating the SCI-induced exacerbated excitatory response to serotonin.

**Figure 4.**
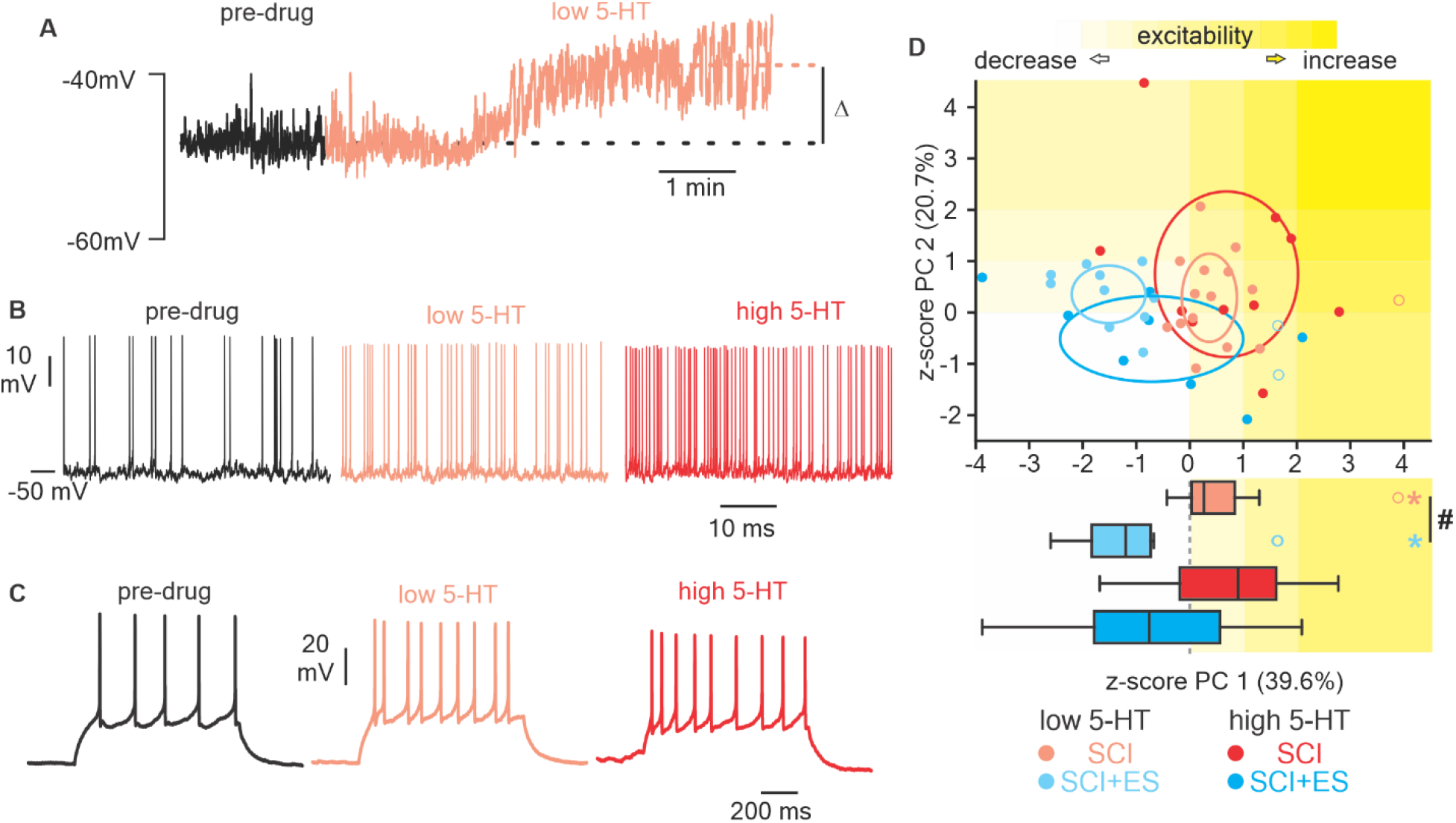
Serotonergic modulation of Shox2 interneurons from SCI and SCI+ES mice. **(A)** Representative trace of effect of low serotonin to the membrane potential of a Shox2 interneuron from a SCI mouse. **(B-C)** Traces from a Shox2 interneuron from a SCI mouse and the effects serotonin on the firing frequency (B) spontaneous and (C) in response to the injection of 1.5 x rheobase. **(D)** First two PCs of the effects of serotonin on Shox2 interneurons from SCI (pink for 0.1 µM serotonin and red for 10 µM serotonin) and SCI+ES (light cyan for 0.1 µM serotonin and cyan for 10 µM serotonin) mice. Ellipses are centered on the mean and semi-axes are generated by the standard error of each two first PCs. Outliers are represented by open circles. The lower compares the z-score of PC1 of the changes in the excitability of Shox2 interneurons. The yellow background indicates an increase in the excitability by serotonin. Low serotonin increases the excitability of Shox2 interneurons from SCI mice (*one-sample *t* test, *p*=0.003) and decreases it from SCI+ES mice (*one-sample *t* test, *p*=0.0001). The response to low serotonin on Shox2 interneurons from SCI and SCI+ES mice was different (# unpaired *t* test, *p*=0.0001).

**Table 2.** Serotonergic modulation of Shox2 interneurons from SCI and SC+ES mice represented as change from baseline (Δ).

| $\Delta$ from baseline | (mean ± SD) | One sample *t* or Wilcoxon test | *p* | (mean ± SD) | One sample *t* or Wilcoxon test | *p* | unpaired *t* or Mann-Whitney test | *p* |
| --- | --- | --- | --- | --- | --- | --- | --- | --- |
| | 0.1 μM 5-HT SCI | | | 0.1 μM 5-HT SCI+ES | | | 0.1 μM 5-HT SCI vs SCI+ES | |
| Resting membrane potential (mV) | 4.5 ± 4.5 | *t*<sub>(14)</sub>=3.9 | *0.002 | -0.4 ± 3.0 | *t*<sub>(11)</sub>=0.4 | 0.7 | *t*<sub>(25)</sub>=3.2 | *0.004 |
| Rheobase (pA) | -0.6 ± 4.9 | *t*<sub>(13)</sub>=0.5 | 0.6 | 11.2 ± 13.6 | *t*<sub>(10)</sub>=2.8 | *0.02 | *t*<sub>(23)</sub>=3.0 | *0.006 |
| Action potential threshold (mV) | -1.9 ± 2.5 | *t*<sub>(14)</sub>=2.9 | *0.01 | 0.3 ± 1.7 | *t*<sub>(11)</sub>=0.6 | 0.5 | *t*<sub>(25)</sub>=2.5 | *0.02 |
| Spontaneous firing frequency (Hz) | 0.15 ± 0.18 | *t*<sub>(13)</sub>=3.1 | *0.008 | -0.05 ± 0.05 | *t*<sub>(9)</sub>=3.1 | *0.01 | *t*<sub>(22)</sub>=3.5 | *0.002 |
| Firing frequency at 1.5x rheobase (Hz) | 1.6 ± 2.7 | *t*<sub>(13)</sub>=2.1 | *0.03 | -3.9 ± 1.5 | *t*<sub>(9)</sub>=8.5 | *0.0001 | *t*<sub>(22)</sub>=5.7 | *0.0001 |
| | 10 μM 5-HT SCI | | | 10 μM 5-HT SCI+ES | | | 10 μM 5-HT SCI vs SCI+ES | |
| Resting membrane potential (mV) | 7.5 ± 10 | *t*<sub>(9)</sub>=2.4 | *0.04 | -1.4 ± 3 | *t*<sub>(7)</sub>=1.3 | 0.2 | *t*<sub>(16)</sub>=2.4 | *0.03 |
| Rheobase (pA) | -1.1 ± 11 | W=-20 | 0.3 | 7.3 ± 33 | W=5 | 0.7 | U=29.5 | 0.4 |
| Action potential threshold (mV) | -1.9 ± 5 | *t*<sub>(9)</sub>=1.2 | 0.3 | -2.9 ± 1.9 | *t*<sub>(7)</sub>=4.1 | 0.004 | *t*<sub>(16)</sub>=0.5 | 0.6 |
| Spontaneous firing frequency (Hz) | 0.4 ± 0.8 | W=23 | 0.2 | 0.1 ± 0.6 | W=-8 | 0.5 | U=24.5 | 0.1 |
| Firing frequency at 1.5x rheobase (Hz) | 2 ± 4.5 | *t*<sub>(9)</sub>=1.4 | 0.2 | -4 ± 5.8 | *t*<sub>(7)</sub>=2.0 | 0.09 | *t*<sub>(16)</sub>=2.5 | *0.02 |

**Table 3.** Comparison of the z-score for the two first principal components for the serotonin modulation of Shox2 interneurons from SCI and SCI+ES mice.

| $\Delta$ from baseline | (mean±SD) | One sample *t* or Wilcoxon test | *p* | (mean ± SD) | One sample *t* or Wilcoxon test | *p* | unpaired *t* or Mann-Whitney test | *p* |
| --- | --- | --- | --- | --- | --- | --- | --- | --- |
| | 0.1 μM 5-HT SCI | | | 0.1 μM 5-HT SCI+ES | | | 0.1 μM 5-HT SCI vs SCI+ES | |
| PC1 (39.6 %) | 0.4 ± 0.5 | $t_{(13)}=2.6$ | *0.02 | -1.5 ± 0.7 | $t_{(9)}=6.8$ | *0.0001 | $t_{(22)}=7.5$ | *0.0001 |
| PC2 (20.7%) | 0.3 ± 0.8 | $t_{(14)}=1.3$ | 0.2 | 0.2 ± 0.7 | $t_{(11)}=0.9$ | 0.4 | $t_{(25)}=0.4$ | 0.7 |
| | 10 μM 5-HT SCI | | | 10 μM 5-HT SCI+ES | | | 10 μM 5-HT SCI vs SCI+ES | |
| PC1 (39.6 %) | 0.7 ± 1.4 | $t_{(9)}=1.7$ | 0.1 | -0.7 ± 1.9 | $t_{(7)}=1.1$ | 0.3 | $t_{(16)}=1.9$ | 0.08 |
| PC2 (20.7%) | 0.3 ± 1.0 | $t_{(8)}=1.0$ | 0.3 | -0.5 ± 0.9 | $t_{(7)}=1.5$ | 0.2 | $t_{(15)}=1.7$ | 0.1 |

**Table 4.** Changes (Δ) in the properties of Shox2 interneurons from SCI and SCI+ES mice after the application of DOI to activate 5-HT_2B/2C_ receptors from the baseline acquired in the presence of the 5-HT_2A_ receptor antagonist, ketanserin.

| $\Delta$ from ketanserin | SCI | | | SCI+ES | | | SCI VS ES | |
| --- | --- | --- | --- | --- | --- | --- | --- | --- |
| | (mean ± SD) | One sample *t* or Wilcoxon test | *p* | (mean ± SD) | One sample *t* or Wilcoxon test | *p* | unpaired *t* or Mann-Whitney test | *p* |
| Resting membrane potential (mV) | 3.0 ± 3.1 | $t_{(10)}=3.3$ | *0.009 | 2.8 ± 1.6 | W=28 | *0.02 | U=37 | 0.9 |
| Rheobase (pA) | -0.5 ± 5.1 | W=-1 | 0.9 | -1.3 ± 2.4 | W=-9 | 0.4 | U=28 | 0.4 |
| Action potential threshold (mV) | -2.1 ± 1.8 | $t_{(10)}=3.9$ | *0.003 | -0.6 ± 1.6 | W=-9 | 0.4 | U=22 | 0.2 |
| Spontaneous firing frequency (Hz) | 0.98 ± 1.5 | W=41 | *0.01 | -1.4 ± 1.5 | W=-28 | *0.02 | U=2 | *0.0002 |
| Firing frequency at 1.5x rheobase (Hz) | 3.5 ± 5.5 | $t_{(10)}=2.1$ | 0.06 | 1.0 ± 4.9 | W=1 | 0.9 | U=25.5 | 0.3 |

**Table 5.**
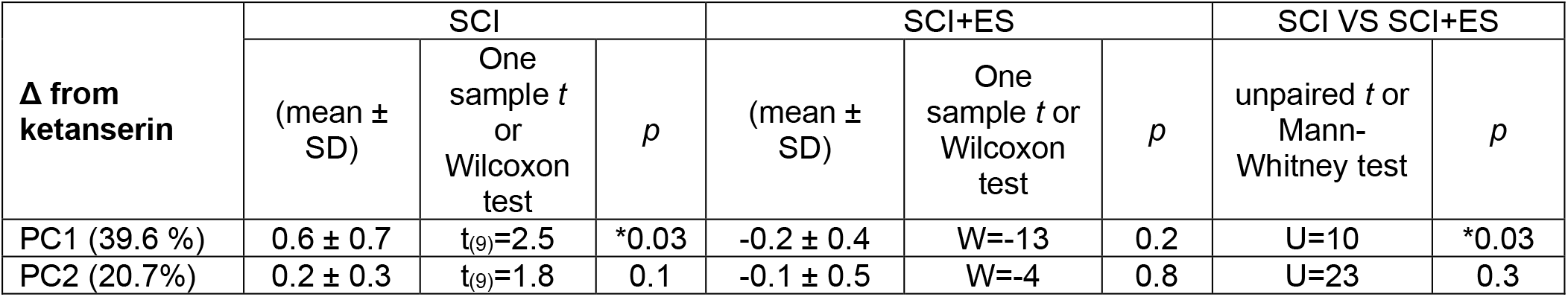
Comparison of the z-score for the two first principal components of the response to the activation of 5-HT_2B/2C_ receptors on Shox2 interneurons from SCI and SCI+ES mice.

**Table 6.**
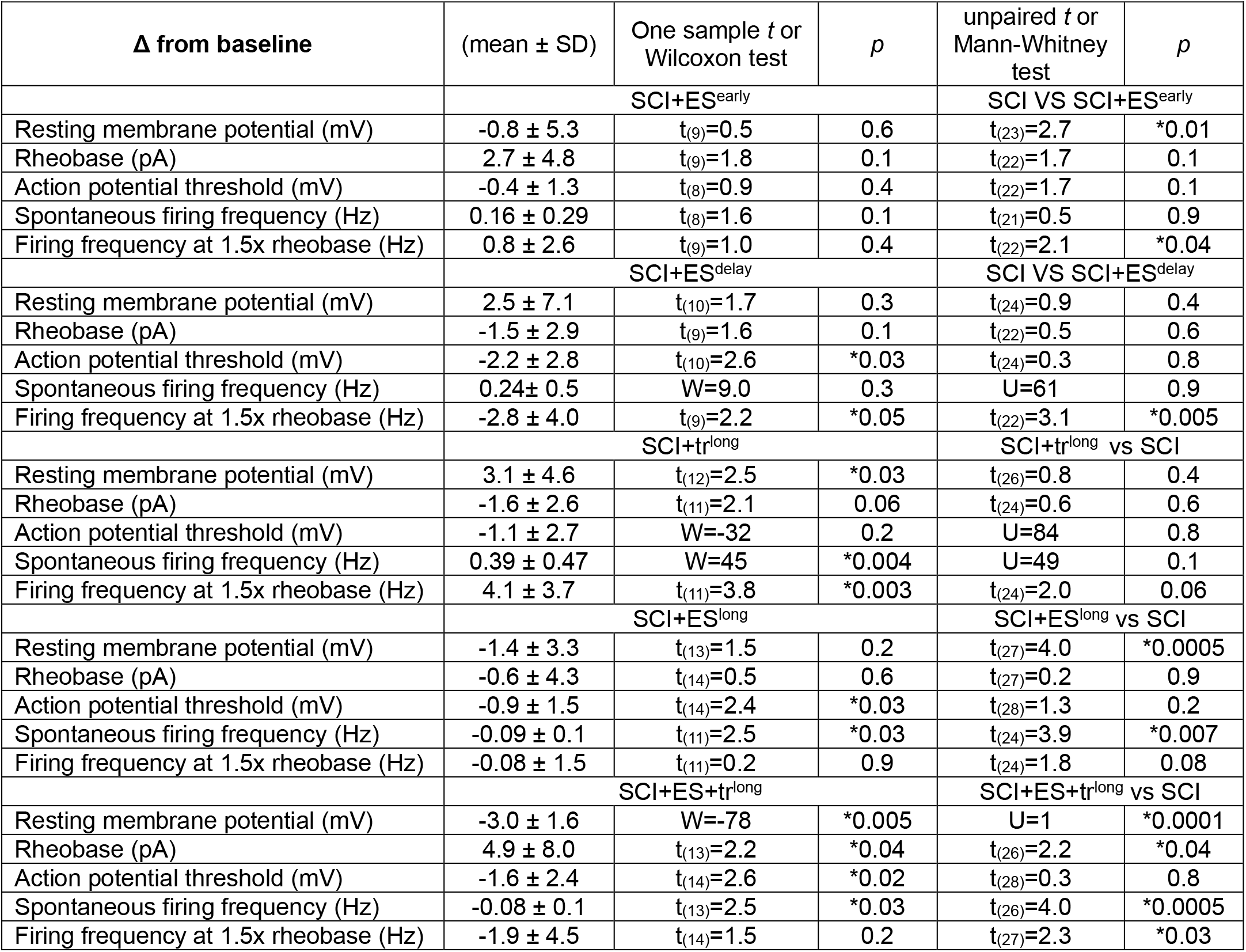
Changes (Δ) in the properties of Shox2 interneurons from SCI+ES^early^, SCI+ES^delay^, SCI+tr^long^ SCI+ES^long^ and SCI+ES+tr^long^ mice after the application of low serotonin concentrations. Comparison to baseline (one sample *t*-test or Wilcoxon test) and to changes in properties of Shox2 interneurons from SCI mice in response to low serotonin concentrations (unpaired *t*-test or Mann-Whitney test).

**Table 7.**
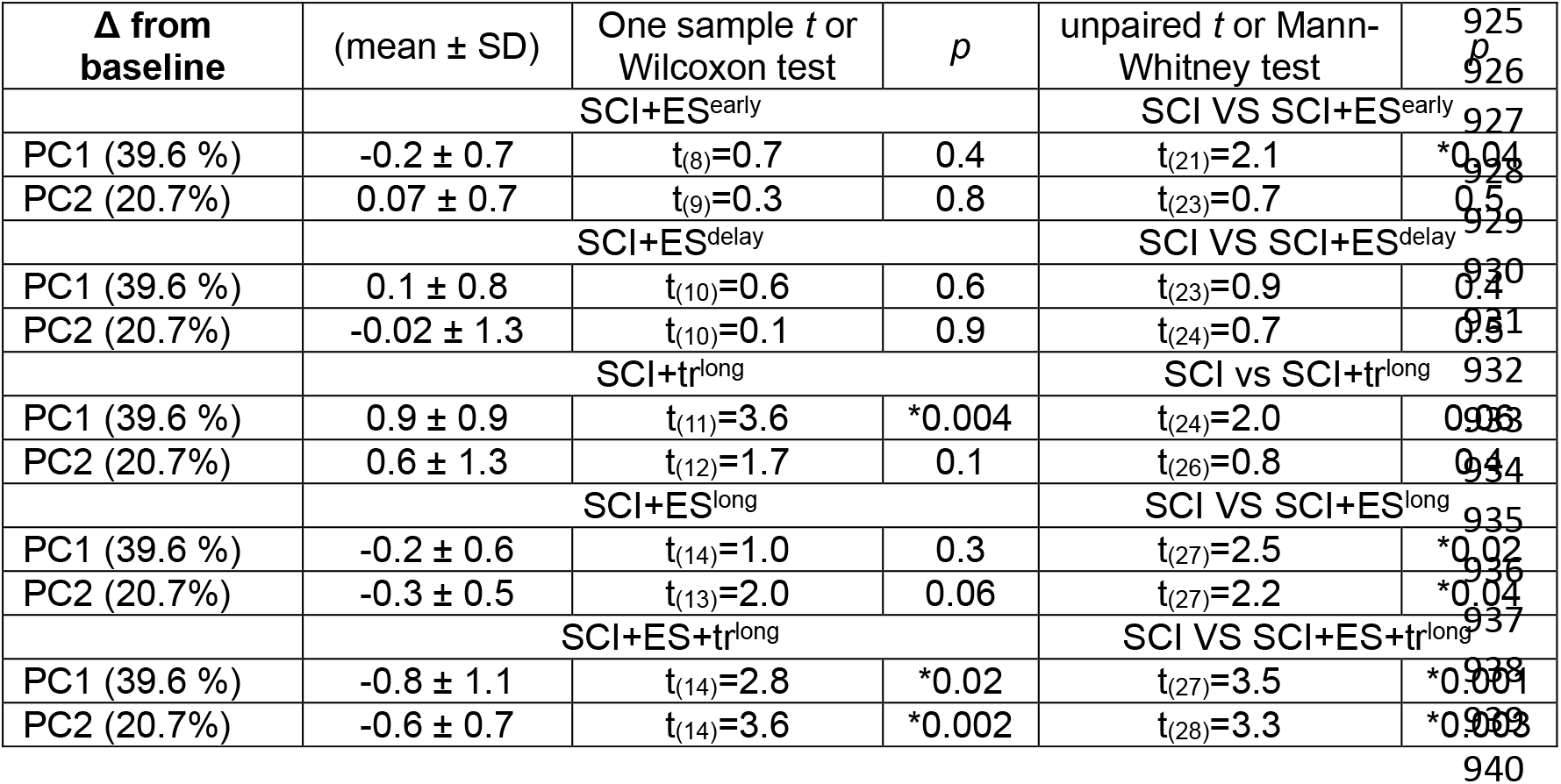
Comparison of the z-score for the two first principal components of the serotonin modulation of Shox2 interneurons from SCI+ES^early^, SCI+ES^delay^, SCI+tr^long^ SCI+ES^long^ and SCI+ES+tr^long^ mice and comparison with SCI mice group.

An increase in the expression or activation of 5-HT_2B/2C_ has been observed in spinal neurons after SCI leading to supersensitivity to serotonin (*25, 27, 43–45*). Here, the lack of excitatory response to serotonin in Shox2 interneurons from SCI+ES mice, led us to hypothesize that activation of 5-HT_2B/2C_ receptors would have less of an effect on Shox2 interneurons from SCI+ES mice compared to SCI mice. We applied a 5-HT_2_ receptor agonist, DOI, together with the specific antagonist for 5-HT_2A_ receptors, ketanserin. We compared the effects produced by the agonist and antagonist combination with the effects produced by antagonist alone on the same electrophysiological parameters and reduced dimensionality with PCs as described above in Shox2 interneurons from SCI and SCI+ES mice (Figure 5A, Table 4 and 5 and supplementary Table 3). We found that the activation of 5-HT_2B/2C_ receptors increased the excitability of Shox2 interneurons from SCI mice (PC1 z-score = 0.6 ± 0.7; one-sample *t* test, *t*(_9_) = 2.6, *p* = 0.03; Figure 5B, light red; Table 5). However, the activation of 5-HT_2B/2C_ receptors did not alter the excitability of Shox2 interneurons from SCI+ES mice (PC1 z-score of −0.2 ± 0.4; Wilcoxon signed rank test, W=-13, *p* = 0.2; Figure 5B, light cyan; Table 5). The response to the activation of 5-HT_2B/2C_ receptors was significantly different between Shox2 interneurons from SCI and SCI+ES mice (Mann-Whitney test, *p* = 0.03; Figure 5B, Table 5). This demonstrates that the 5-HT_2B/2C_ receptors are not present or activatable to modulate Shox2 interneurons following SCI with ES.

**Figure 5.**
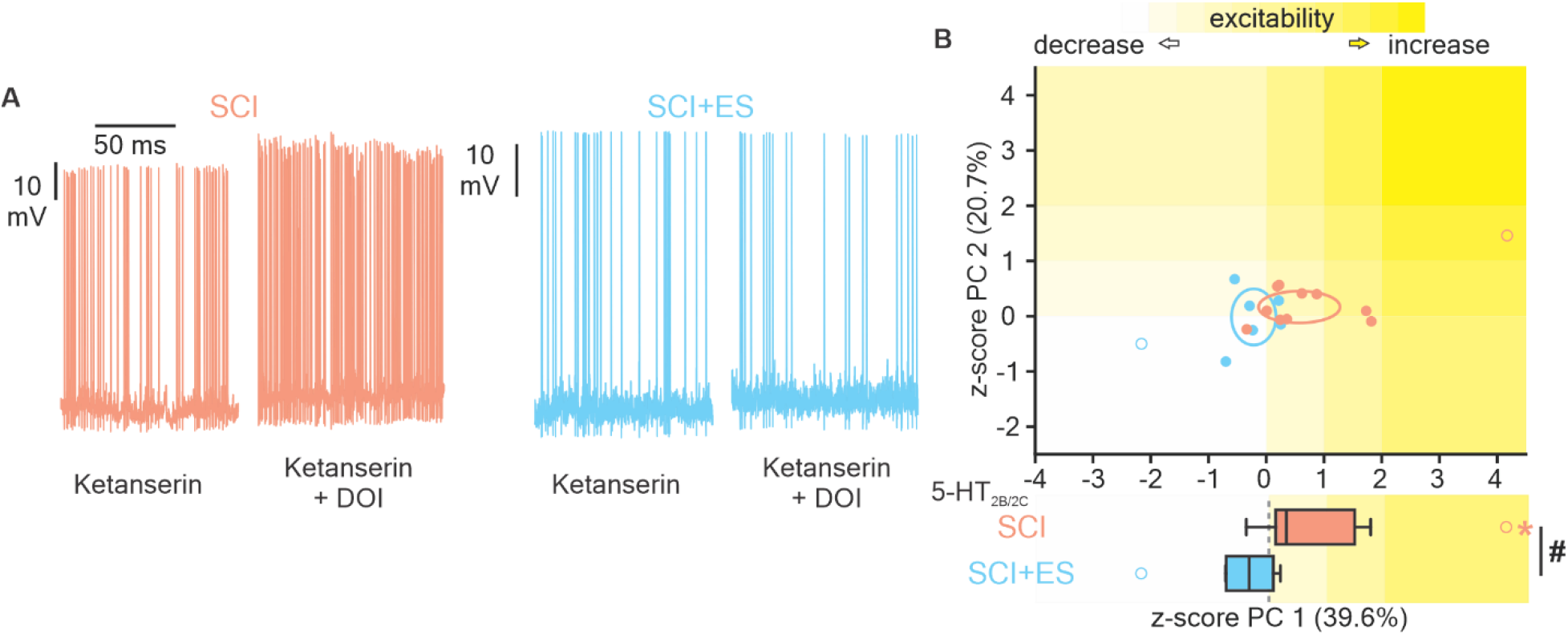
Activation of 5-HT_2B/2C_ receptors on Shox2 interneurons from SCI and SCI+ES mice. **(A)** Representative traces of spontaneous firing of Shox2 interneurons in the presence of the 5-HT_2A_ antagonist, ketanserin (left trace) and ketanserin in combination with the 5-HT_2_ receptor agonist, DOI (right trace) to measure the effects of the activation of 5-HT_2B/2C_ receptors on Shox2 interneurons from SCI (pink) and SCI+ES (cyan) mice. **(B)** Representation of the first two PCs of the changes in the excitability of Shox2 interneurons from SCI (pink) and SCI+ES (light cyan) in response to the activation of 5-HT_2B/2C_ receptors. Ellipses and outliers are represented as in Figure 4. The lower graph compares the z-score of PC1 for the activation of 5-HT_2B/2C_ receptors on Shox2 interneurons from SCI and SCI+ES mice. The yellow background indicates an increase in the excitability of Shox2 interneurons after the activation of 5-HT_2B/2C_ receptors. The activation of 5-HT_2B/2C_ receptors increases the excitability of Shox2 interneurons from SCI mice (*one-sample *t* test, *p* = 0.03) but not from SCI+ES mice (Wilcoxon signed rank test, *p* = 0.09). The response to activation of 5-HT_2B/2C_ receptors on Shox2 interneurons from SCI and SCI+ES mice was different (# Mann-Whitney test, *p* = 0.045).

### ES effects on afferent input and serotonin modulation of Shox2 interneurons persist when ES is discontinued

So far, we showed that the sensory afferent input pathways to Shox2 interneurons are different from mice that received ES in comparison to untreated SCI counterparts (Figure 2). Additionally, the increased excitability of Shox2 interneurons from SCI mice in the presence of low concentrations of serotonin is not observed in Shox2 interneurons from SCI+ES mice (Figure 4). To determine whether the effects of ES on the sensory afferent input pathways and serotonergic modulation of Shox2 interneurons are maintained when ES is discontinued, we included an SCI+ES^early^ group of mice (N=10) which received 3 weeks of ES followed by 4 weeks (4.1 ± 1.1) without treatment (Figure 6A purple). The durations were chosen because 4 weeks of injury and 3 weeks of ES is sufficient for significant differences between SCI and SCI+ES groups (Figures 2 and 4). We found that 60% (6 of 10) of Shox2 interneurons from SCI+ES^early^ mice received only excitatory inputs while 40% (4 of 10 neurons) received inhibitory inputs. Of the inhibitory inputs, half of them were exclusively inhibitory (2 of 10 neurons) and the other half were mixed inhibitory and excitatory (2 of 10 neurons, Figure 6B purple pie graph). The percentage of afferent input types to Shox2 interneurons was significantly different from the SCI mice (*χ*^2^ =13.9, p=0.001) but not the SCI+ES mice (*χ*^2^ =1.1, p=0.6). Additionally, the increase in the excitability of Shox2 interneurons by low serotonin concentrations observed in SCI mice was not present in Shox2 interneurons from SCI+ES^early^ mice (Figure 6C, Table 6 and supplementary Table 1). In fact, the PC1 z-score of the response to low serotonin (−0.8 ± 0.7) was significantly different between Shox2 interneurons from SCI+ES^early^ mice and SCI mice (unpaired *t* test, *t*_(21)_ = 2.1, *p* = 0.04; Table 7). These data suggest that the effects of 3 weeks of early ES are maintained for at least 4 weeks after the discontinuation of ES. Interestingly, the response to low concentrations of serotonin was also significantly different between Shox2 interneurons from SCI+ES^early^ mice and SCI+ES mice (unpaired *t* test, *t*(_15_) = 5.0, p = 0.0002), suggesting that maintained ES may slowly decay with time.

**Figure 6.**
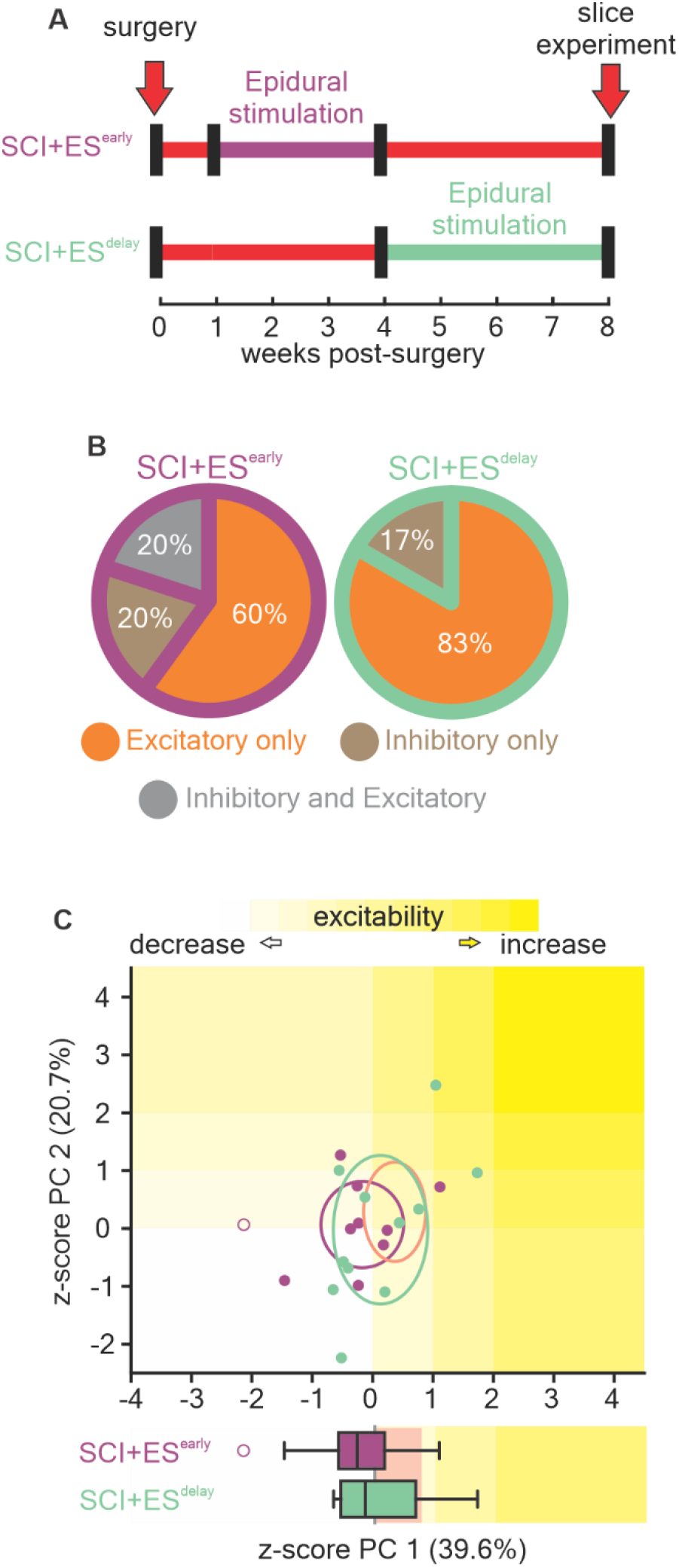
Effects of early and delayed ES on the sensory afferent transmission to Shox2 interneurons and the serotonergic modulation of Shox2 interneurons. **(A)** Experimental timeline of SCI+ES^early^ (purple) and SCI+ES^delay^ (green) mice. **(B)** Incidence of each type of sensory afferent input to Shox2 interneurons from SCI+ES^early^ and SCI+ES^delay^ mice. The types of sensory afferent inputs to Shox2 interneurons are excitatory only (orange), inhibitory only (brown), and mixed excitatory and inhibitory (gray). **(C)** Two-dimensional representation of the first two PCs changes in the excitability of the Shox2 interneurons from SCI+ES^early^ (purple) and SCI+ES^delay^ (green) in response to 0.1 µM serotonin. Ellipses are centered on the mean for the two first PCs and semi-axes are generated by the standard error of each of the two first PCs. Outliers are represented with open circles. Lower graph compares the z-score of PC1 of 0.1 µM serotonin on the excitability of Shox2 interneurons from SCI+ES^early^ (purple box) and SCI+ES^delay^ (green box) mice. The yellow background indicates an increase in the excitability of Shox2 interneurons after serotonin. The pink background indicates the interquartile range of the modulation of Shox2 interneurons from SCI mice by low serotonin concentrations from Figure 4D.

### Delayed ES does not reverse the SCI-induced plasticity

Next, we sought to elucidate whether ES could alter SCI-induced plasticity at the level of Shox2 interneurons if treatment began at more chronic timepoints. Thus, we explored the effects of ES on the SCI-induced plasticity in a group of mice (N=7) that received the SCI surgery 4 weeks prior to the start of ES and then administered ES for at least 3 weeks (4.0 ± 1.6). We referred to this group of mice as SCI+ES^delay^ (Figure 6A green). We found that most Shox2 interneurons from SCI+ES^delay^ mice received only excitatory inputs (83%, 5 of 6) and only one neuron received inhibitory input (17%, 1 of 6, Figure 6B, green pie graph). The percentage of afferent-evoked synaptic input types to Shox2 interneurons was not different from that observed in recordings of Shox2 interneurons from SCI mice (*χ*^2^ = 0.6, p = 0.4) but was different from SCI+ES mice (*χ*^2^ = 10.5, p = 0.005). Further, low serotonin (0.1 µM) did not have an excitatory effect on Shox2 interneurons from SCI+ES^delay^ mice (PC1 z-score = 0.1 ± 0.8, one-sample *t* test, *t*_(10)_ = 0.6, *p* = 0.6; Figure 6C, light green). However the effects of low serotonin on Shox2 interneurons from SCI+ES^delay^ mice was not significand different to what was observed in SCI mice (unpaired *t* test, *t*_(23)_ = 0.9, *p* = 0.4; Table 7); but was significantly different when compared to SCI+ES mice, where low serotonin was inhibitory (unpaired *t* test, *t*_(19)_ = 4.8, p = 0.0001). Although significantly different from SCI+ES mice, the response of Shox2 interneurons from SCI+ES^delay^ mice to low serotonin concentrations was rather variable (Table 6 and Supplementary Table 1), with nearly half (5 of 11) of the Shox2 interneurons displaying higher excitability measures and the other half (6 of 11) displaying a lower excitability. Overall, these data suggest that delayed ES cannot reverse the SCI-induced plasticity at the level of the Shox2 interneurons.

## DISCUSSION

In the present study, we show that the excitatory bias of both sensory afferent input to and serotonergic modulation of Shox2 interneurons seen after SCI is not observed in SCI mice that receive sub-motor threshold ES. Afferent input pathways to Shox2 interneurons and the serotonergic modulation of Shox2 interneurons were similar in mice that received ES for three or more than six weeks, either when allowed to roam/rest freely (SCI+ES or SCI+ES^long^), or while weight supported on a moving treadmill (SCI+ES+tr^long^). Furthermore, the effects of ES persist for at least four weeks post-stimulation. However, if the ES treatment is introduced at more chronic stages of SCI, it is not able to reverse the SCI-induced effects. The present work demonstrates mechanisms by which ES acts on Shox2 interneurons, a specific population of interneurons related to locomotion. Our results reveal that ES, when delivered early, prevents aberrant sensory information effects from reaching locomotor-related interneurons after SCI and affects neuromodulatory sensitivity at the level of the spinal locomotor circuitry, both of which have implications for the treatment of locomotor dysfunctions after SCI.

### Spinal circuit effects of ES in the absence of functional gains

Our goal was to determine mechanisms of action of ES at the level of identified spinal locomotor circuit interneurons. We chose a complete thoracic transection model in which lumbar spinal circuitry is spared due to its distance from the primary injury. This is consistent with the majority of SCIs occurring at cervical and thoracic levels (*46, 47*). Although the majority of SCIs are not anatomically complete (*8, 48*), this allowed for the dissection of purely spinal mechanisms. Additionally, the stimulation parameters used did not result in any noticeable functional improvements, acutely or chronically. Thus, we have a unique model where functional motor gains and related changes in sensory feedback are not contributors to any of the differences observed. Our results demonstrate direct effects of chronic ES on lumbar spinal circuitry, and we interpret our results in that context.

### The effects of SCI and ES on the sensory afferent transmission to the spinal circuitry

Sensory afferents maintain access to spinal locomotor neurons after SCI, but plasticity occurs due to the lack of descending control of both the spinal interneurons and sensory afferent feedback (*6, 13, 14, 19, 35–37, 49, 50*). ES uses sensory afferent fibers to access spinal locomotor circuitry in an attempt to facilitate motor function after SCI (*10, 16, 32, 33, 51–55*). We previously showed that sensory afferent transmission to Shox2 interneurons is both excitatory and inhibitory in uninjured mice (*25, 56*) but changes to mainly excitatory after chronic (>7 weeks) SCI (*25*). Here, we show that afferent input to Shox2 interneurons is almost exclusively excitatory by 4 weeks post-SCI and that daily ES for three weeks prevents the SCI-induced plasticity. The excitability of Shox2 interneurons remains unaltered between uninjured and SCI (*25*) and between SCI and ES. Thus, the plasticity involved must be at the level of the sensory afferent transmission, the interneurons interposed, and/or the synapse between the interposed interneuron and the Shox2 interneuron (Figure 7). There is evidence for SCI-induced plasticity of both sensory afferents (*14, 57–59*) and spinal interneurons (*14, 22, 26, 37, 60–65*), and suggestions of alterations with ES (*66, 67*). ES has recently been reported to reduce neuronal activity in the spinal cord (*28*), which could align with a reduction in the excitability of the excitatory pathways to Shox2 interneurons and possibly others. The percentage of sensory-evoked inhibitory and excitatory inputs to Shox2 interneurons from mice receiving ES is similar to the percentages observed in Shox2 interneurons from uninjured mice (*25*), suggesting that SCI induces disorganized or maladaptive changes (*24, 48, 68*) that are prevented with early ES.

**Figure 7.**
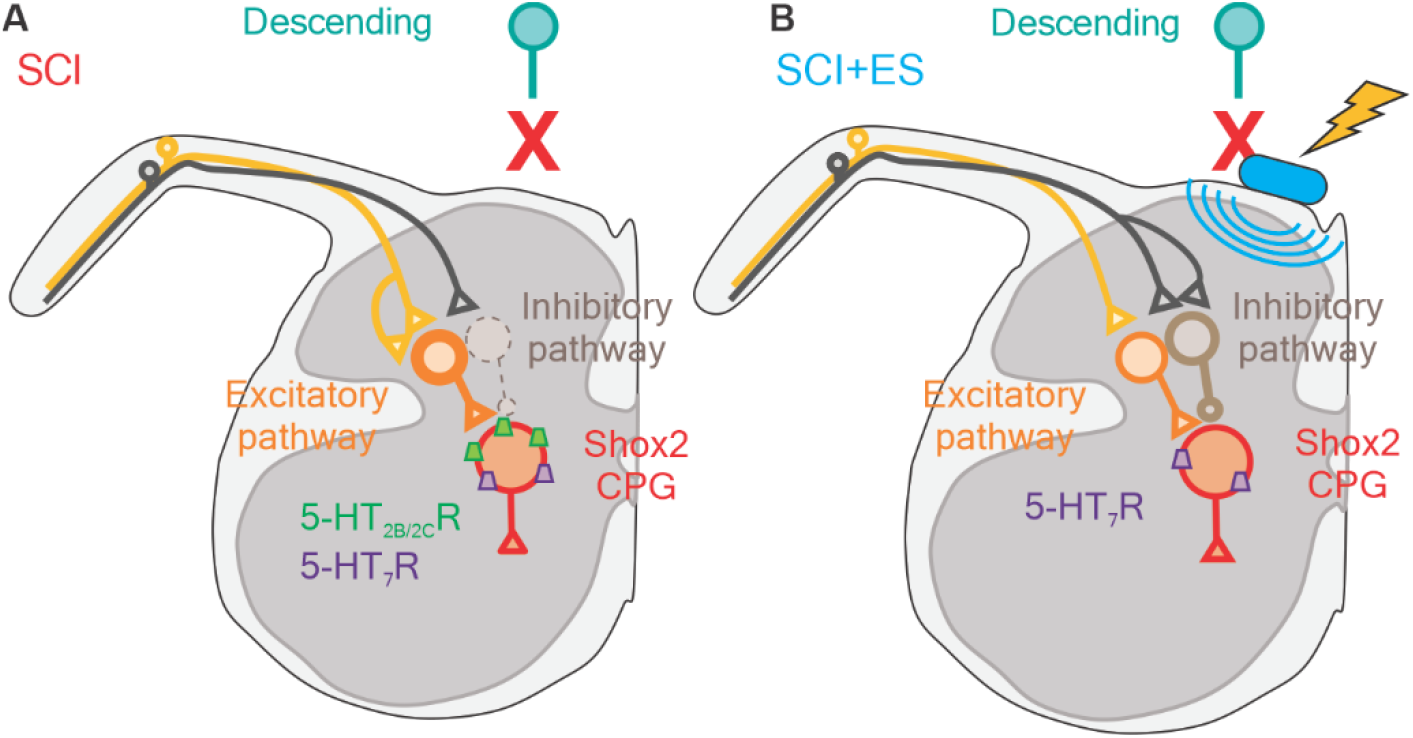
Hypothesis of the ES effects on the SCI-induced plasticity of sensory afferent transmission to and serotonergic modulation of Shox2 interneurons. **(A)** After SCI, the predominant afferent pathway to Shox2 interneurons is excitatory (orange). The serotonergic modulation of Shox2 interneurons is mainly excitatory due to the activation of 5-HT_7_ and 5-HT_2B/2C_ receptors (*25*). **(B)** In mice that received ES, both excitatory and inhibitory sensory afferent pathways to Shox2 interneurons are observed, due to changes in the excitability of excitatory (orange) and/or inhibitory (brown) interneurons interposed between primary afferents and Shox2 interneurons, the afferent inputs to them, or their synaptic connections with Shox2 interneurons. Mice that receive ES treatment do not display the excitatory serotonergic modulation of Shox2 interneurons observed in SCI mice or a response to the activation of 5-HT_2B/2C_ receptors.

### ES affects the serotonergic modulation of Shox2 interneurons

Serotonin in the spinal cord is exclusively released by brainstem descending fibers (*4, 69*) and differentially modulates spinal neurons based on the serotoninergic receptor subtype expressed by the neurons (*69–75*). Serotonin is pro-locomotor, generally inhibiting sensory-related neurons while enhancing motor-related neurons, including Shox2 interneurons (*25, 76–81*). Plasticity of 5-HT receptor expression occurs after SCI leading to serotonin supersensitivity (*25, 27, 39, 41, 43, 44*). Increases in 5-HT_2B/C_ receptor expression contribute to the SCI-induced 5-HT supersensitivity in V2a and Shox2 interneurons (*25, 27*). Although thought to be a homeostatic response to the disruption of the descending serotonergic fibers (*25, 27, 39, 41, 43, 44*), we found that ES prevents serotonin supersensitivity (and related changes in 5-HT_2B/2C_ receptor activation) in Shox2 interneurons. This is an intriguing finding for a complete transection model which should still lack serotonin and suggests that the SCI-induced changes in the spinal serotonergic system are not necessarily due to transmitter levels but may be a more generalized response to the massive reduction in descending activation and consequent altered spinal activation patterns. Further, it suggests that an optimized targeting of cell types by serotonergic drugs, often used experimentally in conjunction with ES (*82–88*), will change over the course of ES treatment.

### Clinical implications

Due to the complexity of the circumstances around a human with a spinal cord injury, clinical use of ES to gain locomotor control often starts late after SCI and yet still with promising outcomes (*89–93*). ES can bring spinal neurons closer to threshold of activation, allowing greater efficacy of descending commands and partial restoration of voluntary control (*16, 94*) and the pairing of afferent and descending input to common spinal neurons strengthens connections (*10, 95*). However, plasticity of the neuronal circuitry below the injury is dynamic (*36, 96, 97*). Our results demonstrate that when ES is delivered early, there are also actions at the level of spinal circuits, stressing the importance of considering the temporal course of ES interventions and possible critical periods for plasticity (*43, 97–100*) and for ES effects. Importantly, the ES strength used here was sub-locomotor threshold and did not result in any obvious concurrent motor activity, making it possible to implement prior to rehabilitation. The lack of significant motor improvements with ES is a strength of our mechanistic study, but it is also a limitation in that we cannot say conclusively that preventing SCI-induced plasticity is functionally beneficial. However, multifaceted (or combination treatments) are likely required (*24, 94, 101*). Future experiments exploring the interactions of early sub-motor threshold ES with spared descending tracts in an incomplete SCI model, in addition to its use prior to suprathreshold ES and/or active rehabilitation, warrant future study.

## METHODS

### Mice

Experiments were performed using the Shox2::Cre; Rosa26-flox-stop-flox-tdTomato (Ai9 from Jax mice, #007909) transgenic mice (*30, 102*). All experimental procedures followed National Institutes of Health guidelines and were approved by the Institutional Animal Care and Use Committee at Drexel University. Mice were randomly assigned to experimental groups prior to surgery. All mice received a complete spinal transection. Mice that were not implanted with ES electrodes formed the SCI or SCI+tr^long^ groups. Mice in ES groups were implanted with ES electrodes and sub-divided depending on the stimulation protocol, as described below.

### SCI and epidural electrodes implantation surgery

Male and female mice 6-10 weeks old (7.9 ± 1.6 weeks) were completely transected at T8/T9 spinal level. Mice were anesthetized with isoflurane (4% induction, 2% maintenance). Dorsal skin was shaved and sterilized with betadine and isopropyl alcohol. To perform the complete spinal cord transection, an incision was performed from the thoracic to lumbar vertebral segments. Following laminectomy, approximately one segment of the thoracic spinal cord was removed at T8-T9. In a subset of transected mice, ES electrodes were implanted during the same surgery.

Two 3-stranded PFA-coated stainless steel wires (A-M systems, #793400) were implanted over the surface of the lumbar spinal segments L2/L3, threaded subcutaneously and mounted to a skull cap. Buprenorphine SR (0.5mg/kg) and either ampicillin (20mg/kg) or Baytril (10mg/kg) were administrated subcutaneously perisurgically. Mice were monitored and bladders were expressed manually twice daily.

### Epidural stimulation parameters

Mice with electrodes implanted received ES by fine ribbon cabling connected to the pin connectors at the skull cap of the mouse on one end and an Isolated Pulse Stimulator (A-M Systems) on the other. The motor threshold of stimulation was tested before the start of each session. ES stimulation was delivered during 10 minutes at 80% threshold (30.9 ± 10.3 µA). In the cases where thresholds were not found but pulses were delivered (as indicated by LED on the stimulator), the stimulation current was 30 μA. ES stimuli were square pulses of 250 μs at 40 Hz.

The SCI+ES mice group received ES for 3 weeks (3.4 ± 0.5 weeks) starting one week after the surgery day, five days per week until one day before the slice experiment. SCI+ES^early^ mice received ES for at least 3 weeks (3.6 ± 0.5 weeks) starting one week after the surgery day, five days per week followed by 4 weeks with no ES (4.1 ± 1.1 weeks). SCI+ES^delay^ mice did not receive ES for 4 weeks (4.3 ± 0.7 weeks) after the surgery, then received ES for at least 3 weeks (4.0 ± 1.6 weeks), five days per week. Both SCI+ES^long^ and SCI+ES+tr^long^ mice groups received ES for more than 6 weeks (7.2 ± 2 and 6.9 ± 1 weeks, respectively) while SCI+tr^long^ mice were on the treadmill with no ES for 8.5 ± 2 weeks starting one week after the surgery day, five days per week until one day before the slice experiment.

### Behavioral testing

Mice were acclimated to the treadmill for 2-3 sessions one week prior to the surgery. The locomotor performance of mice from all groups was tested on the treadmill (5 m/min) in 2 min sessions weekly post-surgery. During these sessions, mice were weight supported over the treadmill with a harness. ES groups were evaluated for 2 min with ES on and 2 minutes with ES off. The locomotor performance was video recorded for off-line analysis. During the analysis, ankle flexions, knee flexions, dorsal steps, and plantar steps were counted and summed. Only mice video recorded on the week of the electrophysiological experiment were included on the locomotor test.

### Spinal slice preparation

For slice experiments, mice were anesthetized with ketamine (150mg/kg) and xylazine (15mg/kg), decapitated, and eviscerated. We visually inspected the lesion site to verify the completeness of the spinal transection. Spinal cords were then removed in ice-cold dissecting solution containing the following (in mM): 222 glycerol, 3 KCl, 11 glucose, 25 NaHCO_3_, 1.3 MgSO_4_, 1.1 KH_2_PO_4_, and 2.5 CaCl_2_ and gassed with 95% O_2_ and 5% CO_2_. The lumbar spinal cord (L2-5) was sectioned transversely (300-350 µm) with dorsal roots attached using a vibrating microtome (Leica Microsystems). Slices were next transferred to an artificial cerebrospinal fluid (ACSF), containing the following (in mM): 111 NaCl, 3 KCl, 11 glucose, 25 NaHCO_3_, 1.3 MgSO_4_, 1.1 KH_2_PO_4_, and 2.5 CaCl_2_ at 37°C for 30min and then passively equilibrated to room temperature for another 30min before recording. Dissecting and recording solutions were continuously aerated with 95%/5% O_2_/CO_2_.

### Whole-cell patch clamp recordings

All recordings were performed at room temperature. Fluorescently labeled (tdTomato) Shox2 interneurons were visualized using a 63x objective lens on a BX51WI scope (Olympus) with LED illumination (Andor Mosaic System) and targeted for whole cell patch clamp recordings. Electrodes were pulled to tip resistances of 5-8 MΩ using a multistage puller (Sutter Instruments) and filled with intracellular solution containing the following (in mM): 128 K-gluconate, 10 HEPES, 0.0001 CaCl2, 1 glucose, 4 NaCl, 5 ATP, and 0.3 GTP. Data were collected with a Multiclamp 700B amplifier (Molecular Devices) and Clampex software (pClamp9, Molecular Devices). Signals were digitized at 20 kHz and filtered at 4 kHz. Glass suction electrodes were used to stimulate dorsal roots as distally as possible. The electrical stimulation was delivered at 0.1Hz frequency with 0.2 ms pulse duration at 100-500 µA in the dorsal root. Mice treated with daily ES mice did not receive ES on the day of the electrophysiology experiment. Neurons with a resting membrane potential more depolarized than −40mV and with an action potential peak that did not reach 0mV were excluded. Active and passive membrane properties were measured as described in Garcia-Ramirez et al. 2022 (*103*).

### Pharmacology

Stock solutions of drugs (1-10 mM) were made and stored at −20°C until needed and then diluted in normal ACSF to experimental concentrations. Ketanserin was dissolved in DMSO resulting in a final DMSO concentration of 0.02% in the bath. Serotonin (5-HT, 0.1 or 10 µM) was bath-applied for 8-10min. In experiments elucidating the participation of specific 5-HT_2B/2C_ receptors, measurements were performed in three conditions: regular ACSF, 8-10 min after the application of the 5-HT_2A_ antagonists ketanserin (1 µM), and 8-10 after a subsequent application of the unspecific 5-HT_2_ agonist DOI (10 µM). All drugs were purchased from Sigma Millipore.

### Experimental design and statistical analyses

Data analysis was performed with Clampfit (Molecular Devices) and MATLAB (The MathWorks). Statistical tests and post hoc analyses used are stated for each experiment and performed with GraphPad Prism (GraphPad Software). Data in Results and Tables are presented as mean ± SD. Statistical significance was set at p<0.05. The distribution of the data was determined by Shapiro–Wilk normality test. The statistical analysis used depended on the test for normal distribution and the experimental conditions. In the case of normally distributed data, we performed paired *t*-tests (comparison between before and after drug conditions), one-sample t tests (comparison between % of control or delta values) and unpaired *t*-tests (comparison between groups of mice). In the cases where data were not normally distributed, we performed Wilcoxon matched pairs test (comparison between before and after drug conditions), Wilcoxon signed rank test (comparison between % of control or deltas) and Mann–Whitney test (comparison between groups of mice). In the experiments considering fraction of total data, we performed binomial tests (for two conditions) and *χ*2 tests (for more than two conditions). Principal component analysis (PCA) was performed to reduce dimensionality of the changes produced by the drugs (serotonin or agonist and antagonist) on the 5 electrophysiological parameters of excitability measured (resting membrane potential, rheobase, action potential threshold, spontaneous firing frequency, and action potential frequency in response a depolarizing current step. The spontaneous firing frequency was measured at approximately −50mV, with the injection of bias current when needed. The step corresponding to 1.5x rheobase was used for the evoked frequency. PCA was performed applying MATLAB algorithms. On figure plots, the dots represent individual data points. For the box plots, the bottom and top of the box represent the 25th and 75th percentiles of the sample (interquartile range), respectively. The line inside the box represents the median. The lines extending above and below each box go from the end of the interquartile range to the furthest observation. Circles are outlier observations, defined as values that were more than 1.5 times the interquartile range away from the bottom or top of the box.

## List of Supplementary Materials

Supplementary Figure 1

Supplementary Figure 2

Supplementary table 1

Supplementary table 2

Supplementary table 3

## Acknowledgements

The authors are grateful to Kendall Schmidt for assistance with initial ES experiments, ULAR staff for expert animal care, and the members of the Dougherty lab and Marion Murray Spinal Cord Research Center for discussions related to the results.

## Author contributions

Conceptualization: DLGR, SFG, KJD

Investigation: DLGR, JRM, NTH, JHW, SJA, LY, NJS, KJD

Visualization: DLGR, KJD Funding acquisition: SFG, KJD

Writing – original draft: DLGR, KJD

Writing – review & editing: DLGR, JRM, NTH, JHW, SJA, LY, NJS, SFG, KJD

## Data and materials availability

All data are available in the main text or the supplementary materials. Further inquiries can be directed to the corresponding author.

---

**Supplementary Figure 1.**
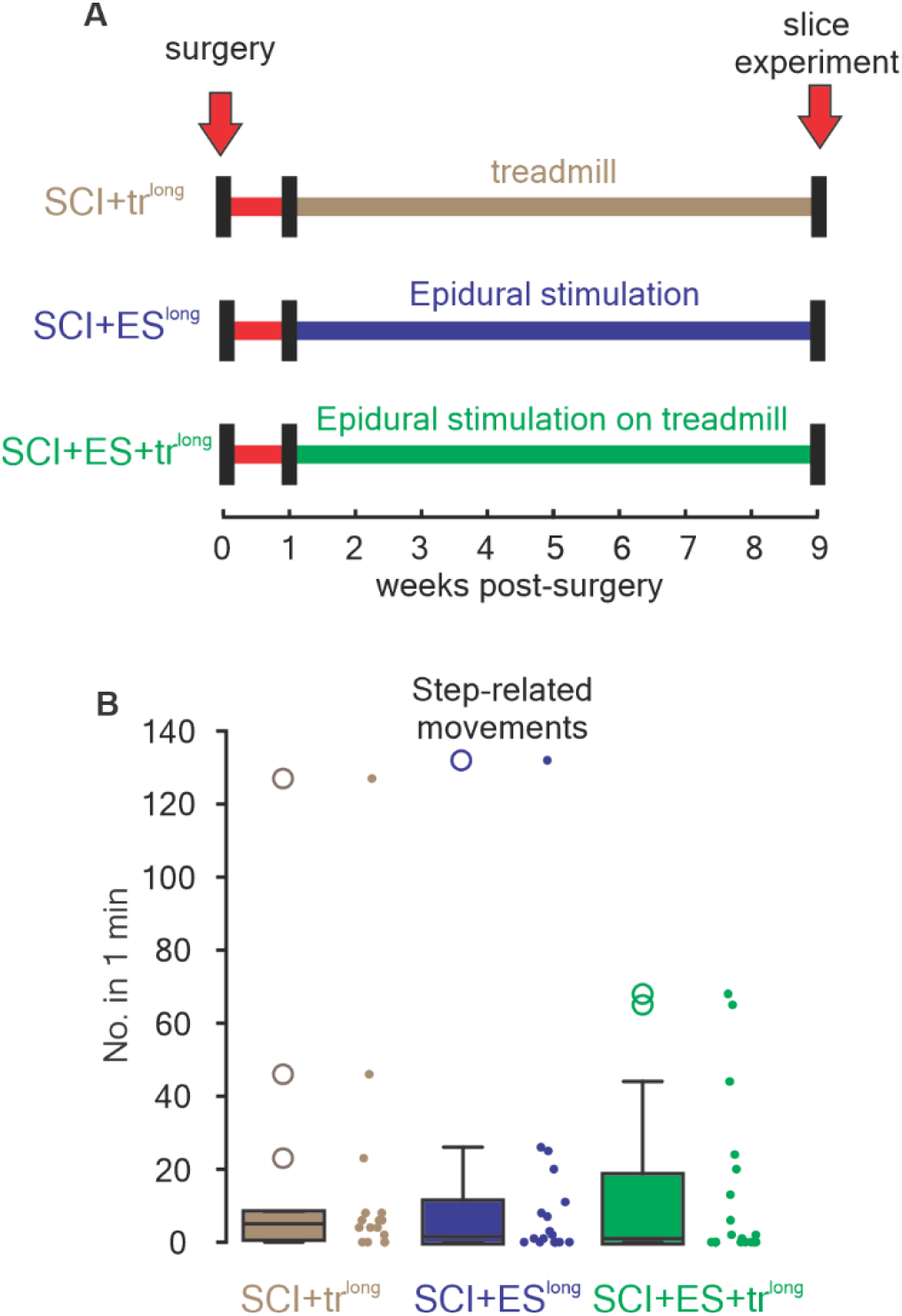
Lengthening ES and delivering ES on the treadmill did not result in locomotor improvements. **(A)** Experimental timeline of SCI+tr^long^ (brown), SCI+ES^long^ (blue) and SCI+ES+tr^long^ (green) mice. **(B)** Comparisons of the number of step-related movements in one minute from SCI+tr^long^ (brown box), SCI+ES^long^ (blue box) and SCI+ES+tr^long^ (green box) mice on the week of the slice experiment.

**Supplementary Figure 2.**
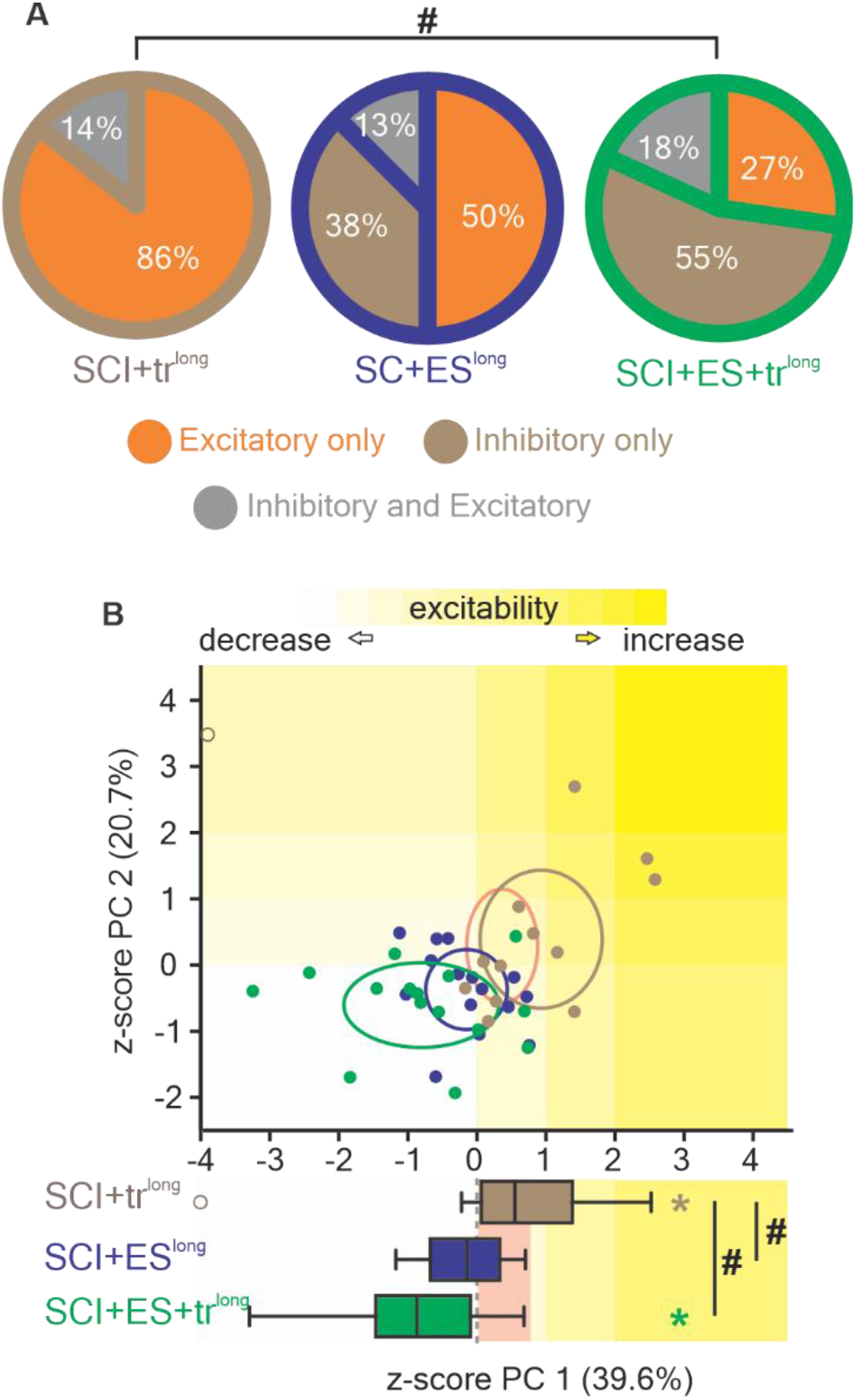
Effects of lengthening ES and delivering ES on the treadmill on the sensory afferent transmission to Shox2 interneurons and the serotonergic modulation of Shox2 interneurons. **(A)** Incidence of each type of sensory afferent input to Shox2 interneurons from SCI+tr^long^ (brown), SCI+ES^long^ (blue) and SCI+ES+tr^long^ (green) mice. The types of sensory afferent inputs to Shox2 neurons are excitatory only (orange), inhibitory only (brown), and mixed excitatory and inhibitory (gray). **(B)** Two-dimensional representation of the first two PCs of the changes in excitability of Shox2 interneurons from SCI+tr^long^ (brown), SCI+ES^long^ (blue) and SCI+ES+tr^long^ (green) mice in response to 0.1 µM. Ellipses are centered on the mean for the two first PCs and semi-axes are generated by the standard error of each of the two first PCs. The lower graph compares the z-score of the PC1 of low serotonin on the excitability of Shox2 interneurons from SCI+tr^long^ (brown), SCI+ES^long^ (blue) and SCI+ES+tr^long^ (green) mice. The yellow background indicates an increase in the excitability of Shox2 interneurons after serotonin. The light red background indicates the interquartile range from the low serotonin modulation of Shox2 interneurons from SCI mice.

**Supplementary table 1.** Properties of Shox2 interneurons from SCI, SCI+ES, SCI+ES^early^, SCI+ES^delay^, SCI+tr^long^, SCI+ES^long^ and SCI+ES+tr^long^ mice after the application of serotonin.

|  | Pre-drug<br>(mean ± SD) | 5-HT<br>(mean ± SD) | paired *t* or<br>Wilcoxon<br>matched<br>pairs test | *p* |
| --- | --- | --- | --- | --- |
| **SCI (low [5-HT])** |  |  |  |  |
| Resting membrane potential (mV) | -55.6 ± 8 | -51.2 ± 9 | *t*<sub>(14)</sub>=3.9 | *0.002 |
| Rheobase (pA) | 27.3 ± 21.3 | 24.8 ± 15.1 | W=-18 | 0.5 |
| Action potential threshold (mV) | -35.8 ± 4.2 | -37.7 ± 5.0 | *t*<sub>(14)</sub>=2.8 | *0.01 |
| Spontaneous firing frequency (Hz) | 0.11 ± 0.2 | 0.35 ± 0.4 | W=60 | *0.005 |
| Firing frequency at 1.5x rheobase (Hz) | 6.6 ± 2.5 | 8.9 ± 3.7 | W=62 | *0.03 |
| **SCI+ES (low [5-HT])** |  |  |  |  |
| Resting membrane potential (mV) | -53.9 ± 9.8 | -54.2 ± 9.2 | *t*<sub>(11)</sub>=0.4 | 0.7 |
| Rheobase (pA) | 21.3 ± 12.7 | 28.3 ± 15.7 | *t*<sub>(11)</sub>=1.3 | 0.2 |
| Action potential threshold (mV) | -37.9 ± 5.3 | 37.8 ± 5.8 | *t*<sub>(11)</sub>=0.6 | 0.6 |
| Spontaneous firing frequency (Hz) | 0.24 ± 0.4 | 0.37 ± 0.7 | W=-11 | 0.6 |
| Firing frequency at 1.5x rheobase (Hz) | 7.2 ± 2.6 | 3.4 ± 4.1 | W=-56 | *0.03 |
| **SCI (high [5-HT])** |  |  |  |  |
| Resting membrane potential (mV) | -54.3 ± 8.7 | -46.8 ± 9.2 | *t*<sub>(9)</sub>=2.4 | *0.04 |
| Rheobase (pA) | 21.2 ± 7.1 | 20.1 ± 12.2 | W=-20 | 0.2 |
| Action potential threshold (mV) | -35.3 ± 4.7 | -37.2 ± 4.9 | *t*<sub>(9)</sub>=1.2 | 0.3 |
| Spontaneous firing frequency (Hz) | 0.16 ± 0.2 | 0.57 ± 0.9 | W=23 | 0.2 |
| Firing frequency at 1.5x rheobase (Hz) | 6.6 ± 2.3 | 8.6 ± 6.0 | *t*<sub>(9)</sub>=1.4 | 0.2 |
| **SCI+ES (high [5-HT])** |  |  |  |  |
| Resting membrane potential (mV) | -50.6 ± 10.1 | -52.0 ± 9.5 | *t*<sub>(7)</sub>=1.3 | 0.2 |
| Rheobase (pA) | 25.0 ± 14.0 | 32.3 ± 28.6 | W=5 | 0.7 |
| Action potential threshold (mV) | -39.8 ± 4.9 | -42.7 ± 4.1 | *t*<sub>(7)</sub>=4.2 | *0.04 |
| Spontaneous firing frequency (Hz) | 0.33 ± 0.5 | 0.42 ± 0.6 | W=-8 | *0.5 |
| Firing frequency at 1.5x rheobase (Hz) | 8.4 ± 2.3 | 4.4 ± 4.8 | W=-22 | 0.2 |
| **SCI+ES<sup>early</sup> (low [5-HT])** |  |  |  |  |
| Resting membrane potential (mV) | -52.7 ± 4.5 | -53.5 ± 6.7 | *t*<sub>(9)</sub>=0.5 | 0.6 |
| Rheobase (pA) | 13.0 ± 7.6 | 15.7 ± 8.3 | W=12 | 0.2 |
| Action potential threshold (mV) | -36.8 ± 2.5 | -36.7 ± 2.7 | *t*<sub>(9)</sub>=0.1 | 0.9 |
| Spontaneous firing frequency (Hz) | 0.31 ± 0.4 | 0.69 ± 0.8 | W=29 | 0.2 |
| Firing frequency at 1.5x rheobase (Hz) | 5.2 ± 1.8 | 4.4 ± 3.3 | W=-12 | 0.5 |
| **SCI+ES<sup>delay</sup> (low [5-HT])** |  |  |  |  |
| Resting membrane potential (mV) | -50.5 ± 8.5 | -48.0 ± 8.1 | *t*<sub>(10)</sub>=1.2 | 0.3 |
| Rheobase (pA) | 22.0 ± 9.7 | 20.7 ± 14.6 | W=-9 | 0.4 |
| Action potential threshold (mV) | -40.1 ± 4.0 | -42.3 ± 4.8 | *t*<sub>(10)</sub>=2.6 | *0.02 |
| Spontaneous firing frequency (Hz) | 0.71 ± 0.9 | 0.97 ± 1.3 | W=12 | 0.4 |
| Firing frequency at 1.5x rheobase (Hz) | 7.2 ± 3.9 | 5.7 ± 5.5 | *t*<sub>(10)</sub>=0.8 | 0.4 |
| **SCI+tr<sup>long</sup> (low [5-HT])** |  |  |  |  |
| Resting membrane potential (mV) | -51.4 ± 5.6 | -48.3 ± 6.3 | *t*<sub>(12)</sub>=2.5 | *0.03 |
| Rheobase (pA) | 25.2 ± 13.3 | 26.8 ± 22.1 | W=-20 | 0.4 |
| Action potential threshold (mV) | -36.3 ± 4.6 | -36.7 ± 3.8 | t<sub>(12)</sub>=0.4 | 0.7 |
| Spontaneous firing frequency (Hz) | 0.2 ± 0.4 | 1.24 ± 1.98 | W=66 | *0.001 |
| Firing frequency at 1.5x rheobase (Hz) | 8.9 ± 5.2 | 11.6 ± 7.2 | t W=66 | *0.03 |
| **SCI+ES<sup>long</sup> (low [5-HT])** |  |  |  |  |
| Resting membrane potential (mV) | -51.2 ± 5.9 | -53.2 ± 6.5 | t<sub>(14)</sub>=1.9 | 0.08 |
| Rheobase (pA) | 25.6 ± 14.9 | 25.0 ± 16.1 | t<sub>(14)</sub>=0.5 | 0.6 |
| Action potential threshold (mV) | -36.6 ± 3.6 | -37.5 ± 4.1 | t<sub>(14)</sub>=2.4 | *0.03 |
| Spontaneous firing frequency (Hz) | 0.35 ± 0.5 | 0.48 ± 0.9 | W=-18 | 0.5 |
| Firing frequency at 1.5x rheobase (Hz) | 6.8 ± 4.4 | 7.0 ± 5.0 | t<sub>(14)</sub>=0.3 | 0.8 |
| **SCI+ES+tr<sup>long</sup> (low [5-HT])** |  |  |  |  |
| Resting membrane potential (mV) | -53.2 ± 6.2 | -56.7 ± 6.2 | t<sub>(14)</sub>=3.3 | *0.005 |
| Rheobase (pA) | 14.1 ± 9.0 | 21.7 ± 18.8 | W=68 | *0.03 |
| Action potential threshold (mV) | -37.0 ± 3.7 | -38.6 ± 4.2 | t<sub>(14)</sub>=2.7 | *0.01 |
| Spontaneous firing frequency (Hz) | 0.16 ± 0.3 | 0.03 ± 0.07 | W=-45 | *0.02 |
| Firing frequency at 1.5x rheobase (Hz) | 6.7 ± 3.4 | 4.9 ± 5.2 | t<sub>(14)</sub>=1.5 | 0.2 |

**Supplementary table 2.** Loading of each variable for the first two principal components.

|  | PC1 | PC2 |
| --- | --- | --- |
| Resting membrane potential (mV) | 0.25 | 0.79 |
| Rheobase (pA) | 0.53 | -0.23 |
| Action potential threshold (mV) | 0.36 | -0.51 |
| Spontaneous firing frequency (Hz) | 0.42 | 0.25 |
| Firing frequency at 1.5x rheobase (Hz) | 0.59 | 0.01 |

**Supplementary table 3.** Properties of Shox2 interneurons from SCI and SCI+ES mice after the activation of 5-HT_2B/2C_ receptors.

|  | Ketanserin<br>(mean ± SD) | Ketanserin + DOI<br>(mean ± SD) | paired *t* or<br>Wilcoxon<br>matched pairs<br>test | *p* |
| --- | --- | --- | --- | --- |
| **SCI** |  |  |  |  |
| Resting membrane potential (mV) | -53.3 ± 5.0 | -50.3 ± 7 | W=58 | *0.007 |
| Rheobase (pA) | 16.1 ± 8.3 | 15.1 ± 8.8 | t<sub>(10)</sub>=0.6 | 0.5 |
| Action potential threshold (mV) | -36.1 ± 6.6 | -38.2 ± 7.2 | t<sub>(10)</sub>=3.9 | *0.003 |
| Spontaneous firing frequency (Hz) | 1.4 ± 0.3 | 2.4 ± 5.7 | W=41 | *0.01 |
| Firing frequency at 1.5x rheobase (Hz) | 5.1 ± 2.0 | 8.6 ± 5.8 | t<sub>(10)</sub>=2.1 | 0.06 |
| **SCI+ES** |  |  |  |  |
| Resting membrane potential (mV) | -53.1 ± 5.5 | -50.3 ± 6.5 | W=28 | *0.02 |
| Rheobase (pA) | 18.4 ± 2.9 | 17.1 ± 3.9 | W=-9 | 0.4 |
| Action potential threshold (mV) | -37.5 ± 3 | -38.1 ± 3.6 | W=-8 | 0.6 |
| Spontaneous firing frequency (Hz) | 2.7 ± 1.8 | 1.3 ± 1.1 | W=-28 | *0.02 |
| Firing frequency at 1.5x rheobase (Hz) | 6.7 ± 2.2 | 7.4 ± 5.4 | W=1 | 0.9 |

